# Biorational Management And Mycosis Studies Of Grape Thrips [*Rhipiphorothrips cruentatus* H.]

**DOI:** 10.1101/2021.06.22.449356

**Authors:** Ankireddy Jawahar reddy, Y. S. Saindane, R. V. Datkhile, B. V. Deore

## Abstract

Field experiment was conducted to test the bioefficacy of various biorational insecticides against grapevine thrips at AICRP on Fruits, Department of Horticulture, MPKV, Rahuri, during 2017-18. Results revealed that standard check emamectin benzoate 5% SG consistently proved to be the most promising by recording the least thrips population (3.10/shoot). Among biorational insecticides neem oil 2% (4.09/shoot) proved as best treatment followed by, karanj oil (4.51/shoot) and neemazol (5.08/shoot). While entomopathogenic fungi *Lecanicillium lecanii* recorded less population (4.24/shoot) and emerged as best treatment as compared to the *Metarhizium anisopliae* (4.87/shoot) and *Beauveria bassiana* (5.34/shoot). However chlli methanolic extract (6.29/shoot), garlic methanolic extract (6.78/shoot), chilli water extract (6.85/shoot) and garlic water extract (7.08/shoot) are the least effective treatments.

Incremental Cost Benefit Ratio (ICBR) in respect of different treatments ranged between 1.30 to 7.92. The highest ICBR of 1:7.92 was recorded in the treatment with emamectin benzoate 5 SG, and it was followed by *Lecanicillium lecanii* (1:6.34) and *Metarhizium anisopliae* (1:5.32). Although neem oil and karanj oil has great reduction of thrips population, but has less incremental cost benefit ratio *i.e.* 2.81 and 3.04, respectively, due to high dose and its cost.

The mycosis test of three entomopathogenic fungi *viz. Beauveria bassiana, Metarhizium anisopliae, Lecanicillium lecanii* were studied on grape thrips. Mycosis by *Beauveria bassiana* was confirmed the pathogenicity of entomopathogenic fungi on grape thrips. Highly pronounced mycosis was observed by *Metarhizium anisopliae* on the dead bodies of thrips. Mycosis test of *Lecanicillium lecanii* was also proved on grape thrips (plate - 1, 2 and 3).

## 1. Introduction

Grapes (*Vitis vinifera* L.) also known as grapevine belonging to the Vitaceae family, originated in Western Asia and Europe. In recent years considerable interest has been aroused in India about grape cultivation due to prolific yield, export potential and good returns. Therefore, the area under grape is constantly increasing. In India, grapes are grown over an area of 1,38,000 ha with the production of 30 lakh MT (Annual Report NRCG, 2017-2018). Maharashtra is the leading grape growing state covering an area of about 78000 ha with the production of 1.80 lakh MT (Annual Report NRCG, 2017-2018).

Thrips, once considered to be the insect pests of minor importance in horticultural crops, but have gained the paramount importance due to their ability to cause economic losses, to subsist on new hosts and by being polyphagous in nature (Dahiya *et al*., 1995). *Ripiphorothrips cruentatus* (H.) and *Scirtothrips dorsalis* (H.) are the species recorded infesting the leaves and berries (Butani., 1979).

Thrips *(Rhipiphoprothrips cruentatus* H.) both nymphs and adults cause damage by rasping the lower surface of the leaf with their stylets and sucking the oozing cell sap. The injured surface is marked by the number of minute spots thereby producing a speckled silvery effect, which can be detected from a distance. They feed in groups, generally on the undersurface of the leaves. Curling of the leaves is observed in case of severe incidence (Kulakarni *et al*., 2007).

Today number of chemicals are often used on large scale regardless of their side effects. Chemical control which also effects the export value of grapes due to pesticide residues. Also some of the results concluded that 27 chemical pesticides out of 171 chemical pesticides can be found usually in grape samples which indicates that the stability of these pesticides is very high or they retain in the grape fruit for a long time after use of them which affect the export value of grapes (Raikwar *et al*., 2011). Hence now a days most of the farmers following biorational pesticide management of thrips which has no residual effects and are ecofriendly. There are number of new biorational pesticides available in the market for the control of various insects. These pesticides are relatively safer, requires in low doses and doesn’t leave residual problems. It is imperative to evaluate such biorational pesticides against thrips, so as to use these insecticides in effective manner or can fit in integrated pest management strategy. With this background, the present study was carried out in an attempt to evaluate the bioefficacy of biorational insecticides and validation of the entomopathegenic fungi growth on grape vine thrips.

## 2. Materials and Methodsa

### 2.1 Bioefficacy of Biorational Insecticides

To evaluate the bioefficacy of certain bio-rational insecticides against thrips on grapevine a field experiment was carried out in a vineyard at All India Co-ordinated Research Project (AICRP) on Fruits, Department of Horticulture, MPKV., Rahuri, after October, 2017 pruning. The grape variety Flame Seedless was chosen for the study. Gardens were selected after ensuring that they were totally unprotected after fruit pruning. The trial was laid out in a Randomized Block Design (RBD) with twelve treatments replicated three times containing two vines each.

#### 2.1.1 Pre and post treatment observations

Pre-treatment count of thrips, was taken prior to the insecticidal application. Eleven insecticides applications were given in the experimental field with the help of a knapsack sprayer. A total of three sprays were applied at an interval of ten days. The data was recorded on population of thrips by tapping five shoots from each treated vines. Observations on thrips and were taken at 3, 5, 7 and 10 days after spray (DAS), (Duraimurugan and Jagadish, 2004). The insecticidal efficacy was assessed by recording the total number of thrips present on vines as well as bunches on two vines in each treatment.

#### 2.1.2 Thrips population assessment

Presence of thrips was recorded on selected shoot and it was expressed as number of thrips per shoot per vine (Kulkarni and Adsule, 2006).

#### 2.1.3 Assessment of effectiveness of insecticides

On the basis of the absolute counts of the thrips recorded, the population reduction in different treatments over control was calculated by using Modified Abbot’s formula given by Fleming and Retnakaran (1985).

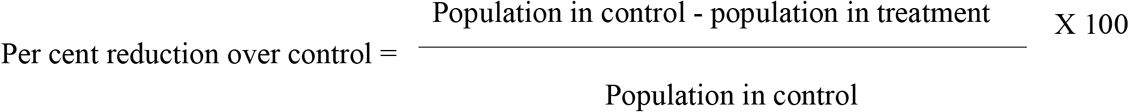

Pre count (1 DBS) and post count (mean of 3, 5, 7 and 10 DAS) population and per cent reduction over control were calculated after each spray. Cumulative mean of three sprays is analysed in order to get the best treatment.

#### 2.1.4 Incremental cost benefit ratio and Yield data

The incremental cost benefit ratio of each insecticide was calculated by taking into account of the prevailing market price of input, produce and labour charges. Grape bunches were harvested from each treatment separately and yield was recorded. Total yield was calculated by adding the yield from different treatments. The per treatment yield was then converted to tonnes per ha.

### 2.2 Mycosis Studies

Three mycoinsecticides like *Beauveria bassiana, Metarhizium anisopliae, Lecanicillium lecanii* were studied for mycosis test on grape thrips. The fungal suspension of three mycoinsecticides are prepared separately in beakers by mixing 5 g of each mycoinsecticide in 100 ml of water in beaker. All the three mycoinsecticides suspensions are prepared in three separate beakers. The young grape leaves are collected from field and their surface is cleaned with mercuric chloride by using cotton, in order to remove fungal spores present on the leaves. Later on the leaves are rinsed with the distilled water to remove the chemical on leaves.

These grape leaves are smeared with the fungal suspension prepared and placed in the petri plates. For each mycoinsecticide three petri plates were prepared for mycosis test. Thrips nymphs were collected from the field and released into each petri plates in numbers of 10. These petri plates were packed with the polythene stripe in order to avoid the escape of thrips from petri plates.These petri plates were incubated in cool place for seven days to promote the infection of fungus on the thrips (Latha *et. al.* 2010).

#### 2.2.1 Method of observation

Detailed microscopic examination of thrips samples collected from the petri plates of different treatments were observed after seven days and ten days of treatment under the stereo microscope with various resolutions like 10 and 40X for the growth of different fungus on various body parts of the thrips. These microscopic photographs are clearly mentioned in the results.

## 3. Results

### 3.1 Bioefficacy of biorational insecticides against grape thrips

The thrips population recorded, a day before spraying (PTC) varied from 8.10 to 8.73 thrips per shoot, which showed non significant difference among treatments indicating homogenous distribution of thrips population in the experimental area (Table 1). There was significant difference among the treatments after 3, 5, 7 and 10 days of first spraying. Considering the mean population of thrips after first spray, it was found that biorational insecticides neem oil (4.21/shoot) and *Lecanicillium lecanii* (4.43/shoot) was the most effective treatment with least population of thrips. Whereas, chilli water extract and garlic water extract was least effective with 7.17 and 7.28 thrips per shoot, respectively. However standard check emamectin benzoate 5 SG @ 11 g a.i.ha^−1^ proved to be significantly superior recording minimum thrips population (3.13/shoot). The data also indicated that higher reduction of population over control was observed in plots treated with standard check emamectin benzoate (64.29%). Among biorational insecticides neem oil (51.90%). *Lecanicillium lecanii* and neemazol had same per cent reduction over control *i.e* 49.33%. Next in order of effectiveness were karanj oil (46.76%), *Metarhizium anisopliae* (44.10%) and *Beauveria bassiana* (38.10%).

**TABLE 1:** Bio-efficacy of some biorational insecticides against thrips on grapes (1^st^ spray)

| Treatment details |  | Dose | Number of thrips per shoot |  |  |  |  | Mean | Per cent reduction of thrips over control |
| --- | --- | --- | --- | --- | --- | --- | --- | --- | --- |
|  |  |  | Precount | 3 DAS | 5 DAS | 7 DAS | 10 DAS |  |  |
| T<sub>1</sub> | *Beauveria bassiana* | 5g/l<br>(1x10<sup>8</sup> cfu/g) | 8.70<br>(3.03) | 8.13<br>(2.94) | 5.70<br>(2.49) | 3.37<br>(1.97) | 4.47<br>(2.23) | 5.42<br>(2.43) | 38.10 |
| T<sub>2</sub> | *Metarhizium anisopliae* | 5g/l<br>(1x10<sup>8</sup> cfu/g) | 8.27<br>(2.96) | 7.43<br>(2.82) | 4.93<br>(2.33) | 3.00<br>(1.87) | 4.20<br>(2.17) | 4.89<br>(2.32) | 44.10 |
| T<sub>3</sub> | *Lecanicillium lecanii* | 5g/l<br>(1x10<sup>8</sup> cfu/g) | 8.10<br>(2.93) | 7.10<br>(2.76) | 4.43<br>(2.22) | 2.30<br>(1.67) | 3.90<br>(2.10) | 4.43<br>(2.22) | 49.33 |
| T<sub>4</sub> | Neem oil (2%) | 20 ml/l | 8.60<br>(3.02) | 7.03<br>(2.74) | 3.63<br>(2.03) | 2.20<br>(1.64) | 3.97<br>(2.11) | 4.21<br>(2.17) | 51.90 |
| T<sub>5</sub> | Karanj oil (2%) | 20 ml/l | 8.50<br>(3.00) | 7.10<br>(2.76) | 3.93<br>(2.11) | 3.07<br>(1.89) | 4.53<br>(2.24) | 4.66<br>(2.27) | 46.76 |
| T<sub>6</sub> | Neemazol (10000ppm) | 3ml/l | 8.10<br>(2.93) | 6.93<br>(2.73) | 3.70<br>(2.05) | 2.87<br>(1.83) | 4.23<br>(2.18) | 4.43<br>(2.22) | 49.33 |
| T<sub>7</sub> | Chilli methanolic extract (2%) | 20 ml/l | 8.17<br>(2.94) | 7.17<br>(2.77) | 5.37<br>(2.42) | 4.30<br>(2.19) | 8.20<br>(2.95) | 6.26<br>(2.60) | 28.48 |
| T<sub>8</sub> | Garlic methanolic extract (2%) | 20 ml/l | 8.27<br>(2.96) | 7.53<br>(2.83) | 5.73<br>(2.50) | 4.90<br>(2.32) | 8.43<br>(2.99) | 6.65<br>(2.67) | 24.00 |
| T<sub>9</sub> | Emamectin benzoate | 0.22g/l | 8.30<br>(2.97) | 5.17<br>(2.38) | 2.10<br>(1.61) | 1.73<br>(1.49) | 3.50<br>(2.00) | 3.13<br>(1.90) | 64.29 |
| T<sub>10</sub> | Chilli water extract (2%) | 20 ml/l | 8.73<br>(3.04) | 8.07<br>(2.93) | 6.47<br>(2.64) | 5.47<br>(2.44) | 8.67<br>(3.03) | 7.17<br>(2.77) | 18.10 |
| T<sub>11</sub> | Garlic water extract (2%) | 20 ml/l | 8.63<br>(3.02) | 8.00<br>(2.92) | 6.60<br>(2.66) | 5.93<br>(2.54) | 8.60<br>(3.02) | 7.28<br>(2.79) | 16.76 |
| T<sub>12</sub> | Untreated Control | --- | 8.67<br>(3.03) | 8.83<br>(3.06) | 8.83<br>(3.06) | 8.60<br>(3.02) | 8.73<br>(3.04) | 8.75<br>(3.04) | 0.00 |
| S. E. m. ± | | --- | 0.117 | 0.088 | 0.075 | 0.065 | 0.097 | 0.061 | |
| CD at 5% |  | --- | NS | 0.260 | 0.220 | 0.193 | 0.286 | 0.181 |  |

\* DAS: Days after spraying; NS: Non significant \* Figures in the parenthesis indicate $\sqrt{x+0.5}$ transformed values

#### 3.1.1 Second Spraying

The results on efficacy of insecticides on population of thrips after second spray were presented in (Table 2).The data on thrips population collected at 10 DAS after I spray was considered as pre count for second spray. Considering the mean population of thrips after second spray, it was found that standard check emamectin benzoate was the most effective treatment with least population of thrips (3.06/shoot). Among biorational insecticides neem oil (4.07/shoot) and *Lecanicillium lecanii* (4.16/shoot) proved as effective treatments. Whereas, chilli water extract and garlic water extract was least effective with 6.85 and 7.10 thrips per shoot, respectively. The cumulative effect of treatments indicated that higher reduction of population over control was observed in plots treated with standard check emamectin benzoate (65.11%). Among biorational imsecticides neem oil (53.61%) emerged as best treatment over control. Next in order of effectiveness were *Lecanicillium lecanii* (52.57%), karanj oil (49.05%), *Metarhizium anisopliae* (43.54%) and *Beauveria bassiana* (37.45%).

**TABLE 2:** Bio-efficacy of some biorational insecticides against thrips on grapes (2^nd^ spray)

| Treatment details |  | Dose | Number of thrips per shoot |  |  |  | Mean | Per cent reduction of thrips over control |
| --- | --- | --- | --- | --- | --- | --- | --- | --- |
|  |  |  | 3 DAS | 5 DAS | 7 DAS | 10 DAS |  |  |
| T<sub>1</sub> | *Beauveria bassiana* | 5g/l<br>(1x10<sup>8</sup> cfu/g) | 8.06<br>(3.02) | 6.00<br>(2.55) | 3.27<br>(1.94) | 4.07<br>(2.14) | 5.48<br>(2.45) | 37.45 |
| T<sub>2</sub> | *Metarhizium anisopliae* | 5g/l<br>(1x10<sup>8</sup> cfu/g) | 8.03<br>(2.92) | 5.47<br>(2.44) | 2.80<br>(1.82) | 3.50<br>(2.00) | 4.95<br>(2.33) | 43.54 |
| T<sub>3</sub> | *Lecanicillium lecanii* | 5g/l<br>(1x10<sup>8</sup> cfu/g) | 7.20<br>(2.77) | 4.83<br>(2.31) | 1.97<br>(1.57) | 2.63<br>(1.77) | 4.16<br>(2.16) | 52.57 |
| T<sub>4</sub> | Neem oil (2%) | 20 ml/l | 6.70<br>(2.68) | 4.67<br>(2.27) | 1.77<br>(1.51) | 3.13<br>(1.91) | 4.07<br>(2.14) | 53.61 |
| T<sub>5</sub> | Karanj oil (2%) | 20 ml/l | 5.93<br>(2.54) | 4.87<br>(2.32) | 3.27<br>(1.94) | 3.80<br>(2.07) | 4.47<br>(2.23) | 49.05 |
| T<sub>6</sub> | Neemazol (10000ppm) | 3ml/l | 7.23<br>(2.78) | 6.00<br>(2.55) | 5.00<br>(2.35) | 4.80<br>(2.30) | 5.76<br>(2.50) | 34.32 |
| T<sub>7</sub> | Chilli methanolic extract (2%) | 20 ml/l | 7.40<br>(2.81) | 6.53<br>(2.65) | 5.03<br>(2.35) | 6.27<br>(2.60) | 6.31<br>(2.61) | 28.04 |
| T<sub>8</sub> | Garlic methanolic extract (2%) | 20 ml/l | 8.10<br>(2.93) | 7.07<br>(2.75) | 6.07<br>(2.56) | 6.97<br>(2.73) | 7.05<br>(2.75) | 19.58 |
| T<sub>9</sub> | Emamectin benzoate | 0.22g/l | 4.30<br>(2.19) | 3.67<br>(2.04) | 1.63<br>(1.46) | 2.63<br>(1.77) | 3.06<br>(1.89) | 65.11 |
| T<sub>10</sub> | Chilli water extract (2%) | 20 ml/l | 7.80<br>(2.88) | 6.73<br>(2.69) | 5.67<br>(2.48) | 7.20<br>(2.77) | 6.85<br>(2.71) | 21.86 |
| T<sub>11</sub> | Garlic water extract (2%) | 20 ml/l | 7.40<br>(2.81) | 7.07<br>(2.75) | 6.33<br>(2.61) | 7.60<br>(2.85) | 7.10<br>(2.76) | 19.01 |
| T<sub>12</sub> | Untreated Control | --- | 8.63<br>(3.02) | 8.67<br>(3.03) | 8.87<br>(3.06) | 8.90<br>(3.07) | 8.77<br>(3.04) | 0.00 |
| S. E. m. ± |  | --- | 0.100 | 0.082 | 0.067 | 0.074 | 0.060 |  |
| CD at 5% |  | --- | 0.295 | 0.241 | 0.197 | 0.217 | 0.176 |  |

\* DAS: Days after spraying; NS: Non significant \* Figures in the parenthesis indicate $\sqrt{x+0.5}$ transformed values.

#### 3.1.2 Third Spraying

The results with regard to the efficacy of treatments after third spray were presented in (Table 3). Considering the mean population of thrips after third spray, it was found that standard check emamectin benzoate was the most effective treatment with least population of thrips (3.11/shoot). Among biorational insecticides neem oil (4.00/shoot), *Lecanicillium lecanii* (4.12/shoot) and karanj oil (4.42/shoot) were the best treatments. Whereas, garlic methanolic extract and garlic water extract was least effective with 6.64 and 6.85 thrips per shoot respectively. The cumulative effect of treatments indicated that higher reduction of population over control was observed in plots treated with standard check emamectin benzoate (63.18%). Among biorational insecticides neem oil is the best treatment with 52.62% reduction over control. Next in order of effectiveness were *Lecanicillium lecanii* (51.23%), karanj oil (47.68%), *Metarhizium anisopilae* (43.63%), neemazol (40.08%) and *Beauveria bassiana* (39.19%).

**TABLE 3:** Bio-efficacy of some biorational insecticides against thrips on grapes (3^rd^ spray)

| Treatment details |  | Dose | Number of thrips per shoot |  |  |  | Mean | Per cent reduction of thrips over control |
| --- | --- | --- | --- | --- | --- | --- | --- | --- |
|  |  |  | 3 DAS | 5 DAS | 7 DAS | 10 DAS |  |  |
| T<sub>1</sub> | *Beauveria bassiana* | 5g/l<br>(1x10<sup>8</sup> cfu/g) | 8.20<br>(2.95) | 5.17<br>(2.38) | 3.13<br>(1.91) | 4.03<br>(2.13) | 5.13<br>(2.37) | 39.19 |
| T<sub>2</sub> | *Metarhizium anisopliae* | 5g/l<br>(1x10<sup>8</sup> cfu/g) | 7.63<br>(2.85) | 5.20<br>(2.39) | 2.80<br>(1.82) | 3.40<br>(1.97) | 4.76<br>(2.29) | 43.63 |
| T<sub>3</sub> | *Lecanicillium lecanii* | 5g/l<br>(1x10<sup>8</sup> cfu/g) | 7.17<br>(2.77) | 4.90<br>(2.32) | 2.10<br>(1.61) | 2.30<br>(1.67) | 4.12<br>(2.15) | 51.23 |
| T<sub>4</sub> | Neem oil (2%) | 20 ml/l | 6.40<br>(2.63) | 4.73<br>(2.29) | 1.97<br>(1.57) | 2.90<br>(1.84) | 4.00<br>(2.12) | 52.62 |
| T<sub>5</sub> | Karanj oil (2%) | 20 ml/l | 6.13<br>(2.58) | 4.80<br>(2.30) | 2.93<br>(1.85) | 3.80<br>(2.07) | 4.42<br>(2.22) | 47.68 |
| T<sub>6</sub> | Neemazol (10000 ppm) | 3ml/l | 7.23<br>(2.78) | 4.80<br>(2.30) | 3.10<br>(1.90) | 5.10<br>(2.37) | 5.06<br>(2.36) | 40.08 |
| T<sub>7</sub> | Chilli methanolic extract (2%) | 20 ml/l | 8.10<br>(2.93) | 6.00<br>(2.55) | 4.80<br>(2.30) | 6.27<br>(2.60) | 6.29<br>(2.61) | 25.47 |
| T<sub>8</sub> | Garlic methanolic extract (2%) | 20 ml/l | 8.27<br>(2.96) | 6.23<br>(2.59) | 5.10<br>(2.37) | 6.97<br>(2.73) | 6.64<br>(2.67) | 21.32 |
| T<sub>9</sub> | Emamectin benzoate | 0.22g/l | 4.40<br>(2.21) | 3.77<br>(2.07) | 1.73<br>(1.49) | 2.53<br>(1.74) | 3.11<br>(1.90) | 63.18 |
| T<sub>10</sub> | Chilli water extract (2%) | 20 ml/l | 7.60<br>(2.85) | 6.23<br>(2.59) | 5.20<br>(2.39) | 7.10<br>(2.76) | 6.53<br>(2.65) | 22.61 |
| T<sub>11</sub> | Garlic water extract (2%) | 20 ml/l | 7.80<br>(2.98) | 6.73<br>(2.99) | 5.67<br>(2.99) | 7.20<br>(3.00) | 6.85<br>(2.99) | 18.85 |
| T<sub>12</sub> | Untreated Control | --- | 8.37<br>(2.88) | 8.43<br>(2.69) | 8.47<br>(2.48) | 8.50<br>(2.77) | 8.44<br>(2.71) | 0.00 |
| S. E. m. ± |  | --- | 0.097 | 0.084 | 0.072 | 0.093 | 0.063 | --- |
| CD at 5% |  | --- | 0.286 | 0.249 | 0.214 | 0.274 | 0.186 | --- |

\* DAS: Days after spraying; NS: Non significant \* Figures in the parenthesis indicate $\sqrt{x+0.5}$ transformed values.

#### 3.1.3 Pooled data

The data pertaining to efficacy of insecticides against thrips during first, second and third spray are pooled and presented in (Table 4). It could be seen that all the insecticidal treatments were significantly superior over untreated control. The pooled data of three sprays revealed that standard check emamectin benzoate 5 SG consistently proved to be the most promising by recording the least population (3.10/shoot). Among biorational insecticides neem oil 2% (4.09/shoot), karanj oil (4.51/shoot) and neemazol (5.08/shoot). While entomopathogenic fungi *Lecanicillium lecanii* recorded less population (4.24/shoot) as compared to the *Metarhizium anisopliae* (4.87/shoot) and *Beauveria bassiana* (5.34/shoot). The data also indicated that higher per cent reduction over control of population was observed in plots treated with standard check emamectin benzoate 5 SG (64.21%). Among biorational insecticides neem oil (52.71%) and *Lecanicillium lecanii* (51.04%). Next in order of effectiveness were karanj oil (47.83%), *Metarhizium anisopliae* (43.76%), neemazol (41.26%), *Beauveria bassiana* (38.23%), chlli methanolic extract (27.35%), garlic methanolic extract (21.63%), chilli water extract (20.83%) and garlic water extract (18.20%) presented in Fig.1.

**TABLE 4:**
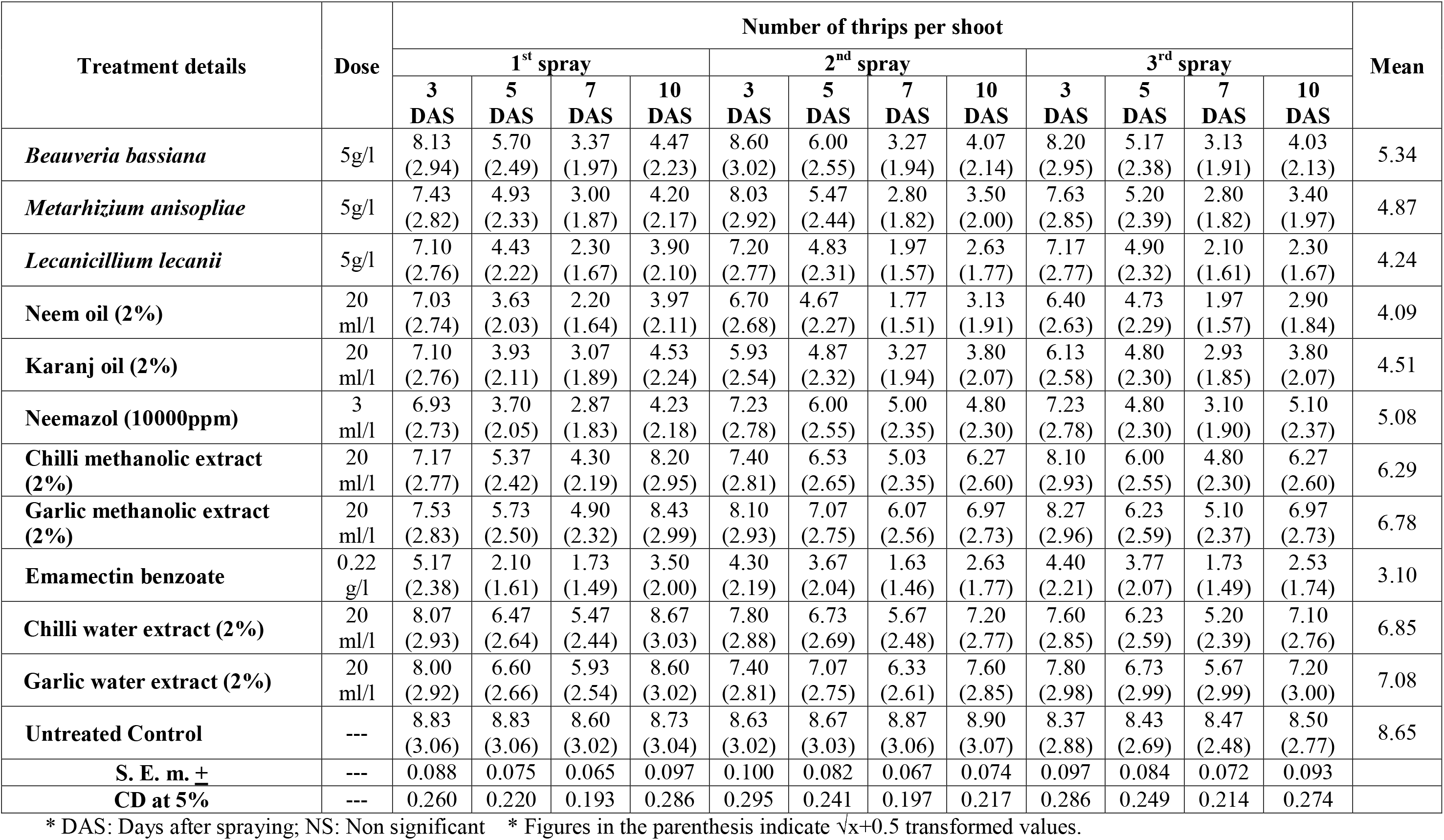
Bio-efficacy of some biorational insecticides against thrips on grapes (pooled data)

**Fig. 1:**
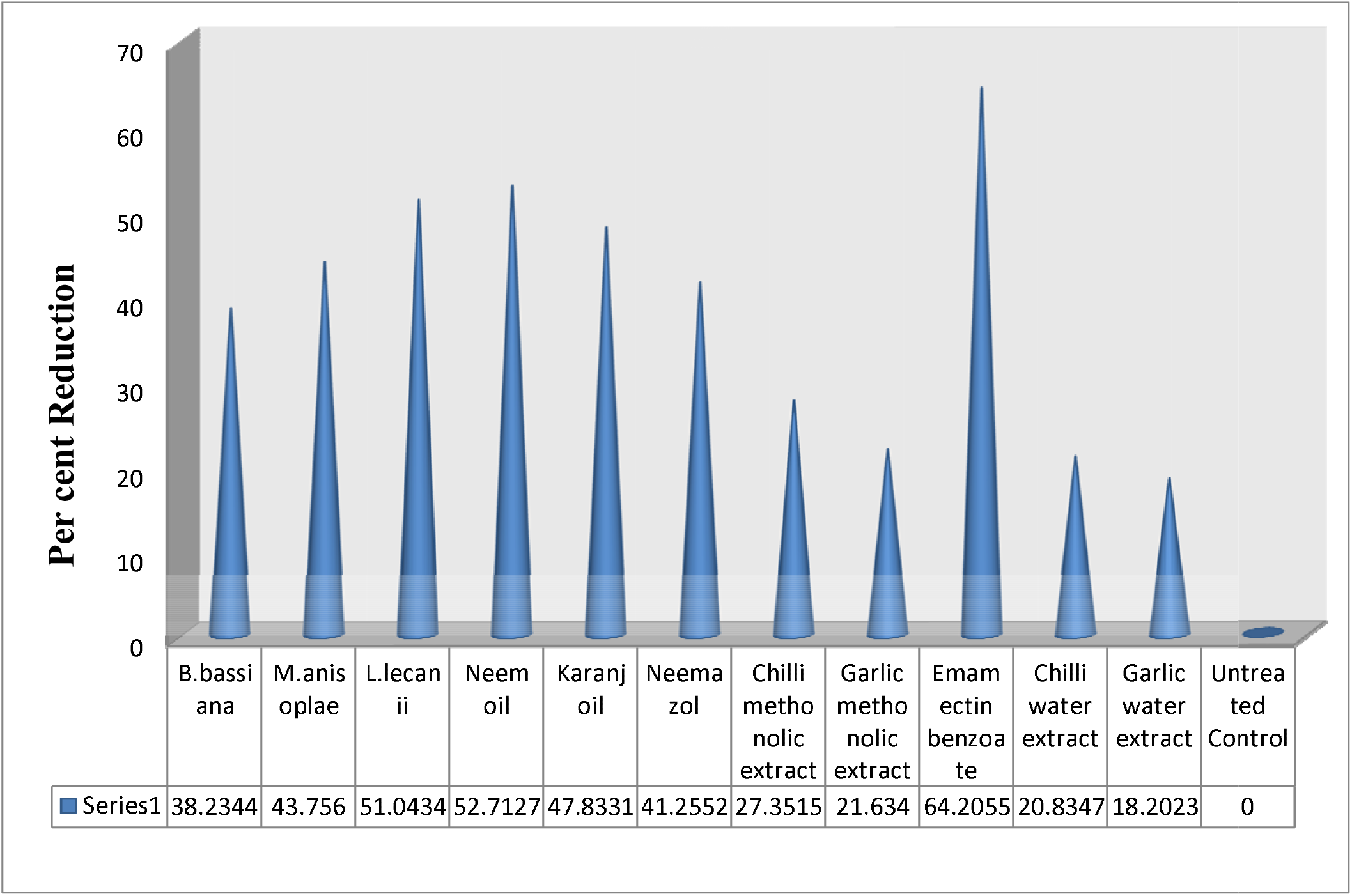
Per cent Reduction over Control of biorational insecticides against thrips on grapes (pooled data)

#### 3.1.4 Cost economics of grapes

The cost effectiveness of the different insecticides used during study was assessed and presented in the (Table 5). The ICBR in respect of different treatments ranged between 1.30 to 7.92. The highest C:B ratio in *L. lecanii* (1:6.34) and *M. anisopliae* (1:5.32). Although neem oil and karanj oil has great reduction of thrips population, but has less cost benefit ratio *i.e* 2.81 and 3.04, respectively due to high cost of the insecticide.

**Table 5:**
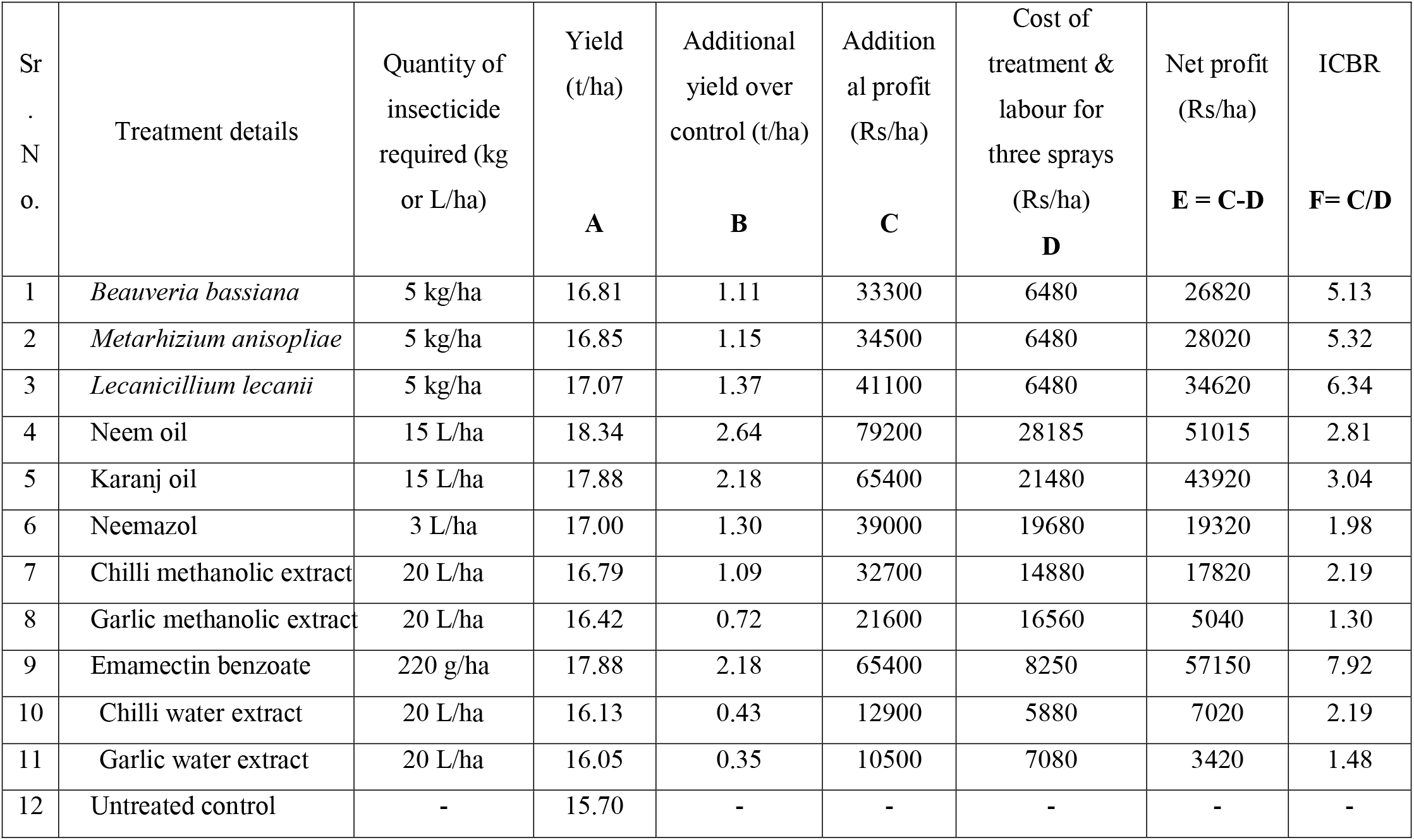
Cost economics of grape production influenced by treatment of biorational insecticides for thrips management.

### 3.2 Mycosis test of mycoinsecticides on grape vine thrips

The fungal suspension of three mycoinsecticides *viz. Beauveria bassiana, Metarhizium anisopliae, Lecanicillium lecanii* were studied for mycosis test on grape thrips. Detailed microscopic examination of the thrips samples collected from the petri plates showed that all the test entomopathogenic fungi found growing in the body of the thrips. The moribund adult thrips showed profuse fungal growth in the body cavity, (Plates 1, 2 and 3). The microscopic photographs in the plates clearly indicated the mycosis by *Beauveria bassiana* was predominant behind compound eyes, prothorax, near fore coxa, stomach portion and between inter segmental spaces (Plate 1 – a). Close up view showed clear growth of fungus in thorax and abdominal portion (Plate 1 – c). In advanced stages after tight filling the body cavity the fungus outgrowth was observed on head, legs and posterior part of abdomen, (Plate 1 – b). Highly pronounced mycosis by *Metarhizium anisopliae* was observed in the thrips which shrunken and hardened its body (Plate 2 – a, b, c and d). The growth was observed in almost all body parts. The growth was observed along inter segmental joints around genital parts (Plate 2 – c and d), tergo-sternum joint (Plate 2 – b), head and prothorax and inter wings (Plate 2 – a and c). *Lecanicillium lecanii* soften the body of the thrips and growth was observed on antennary tips, around compound eyes, legs and tissues in different part of the body and alimentary canal in mid infestation (Plate 3 – a), in advanced stages whole body was captured by the fungus and growth was also vivid on surface of the body (Plate 3 – b and c).

**Plate 1:**
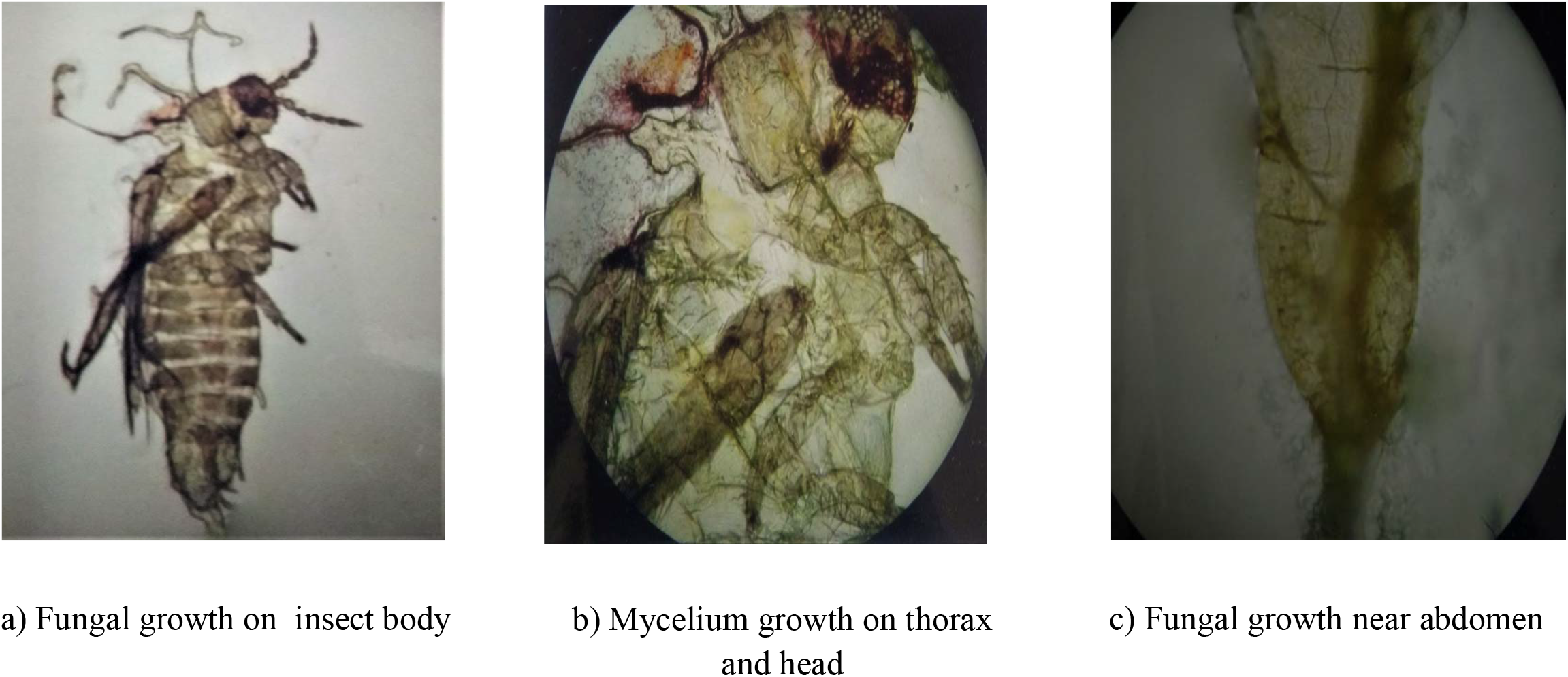
Mycosis of *B.eauveria bassiana* on grape thrips.

**Plate 2:**
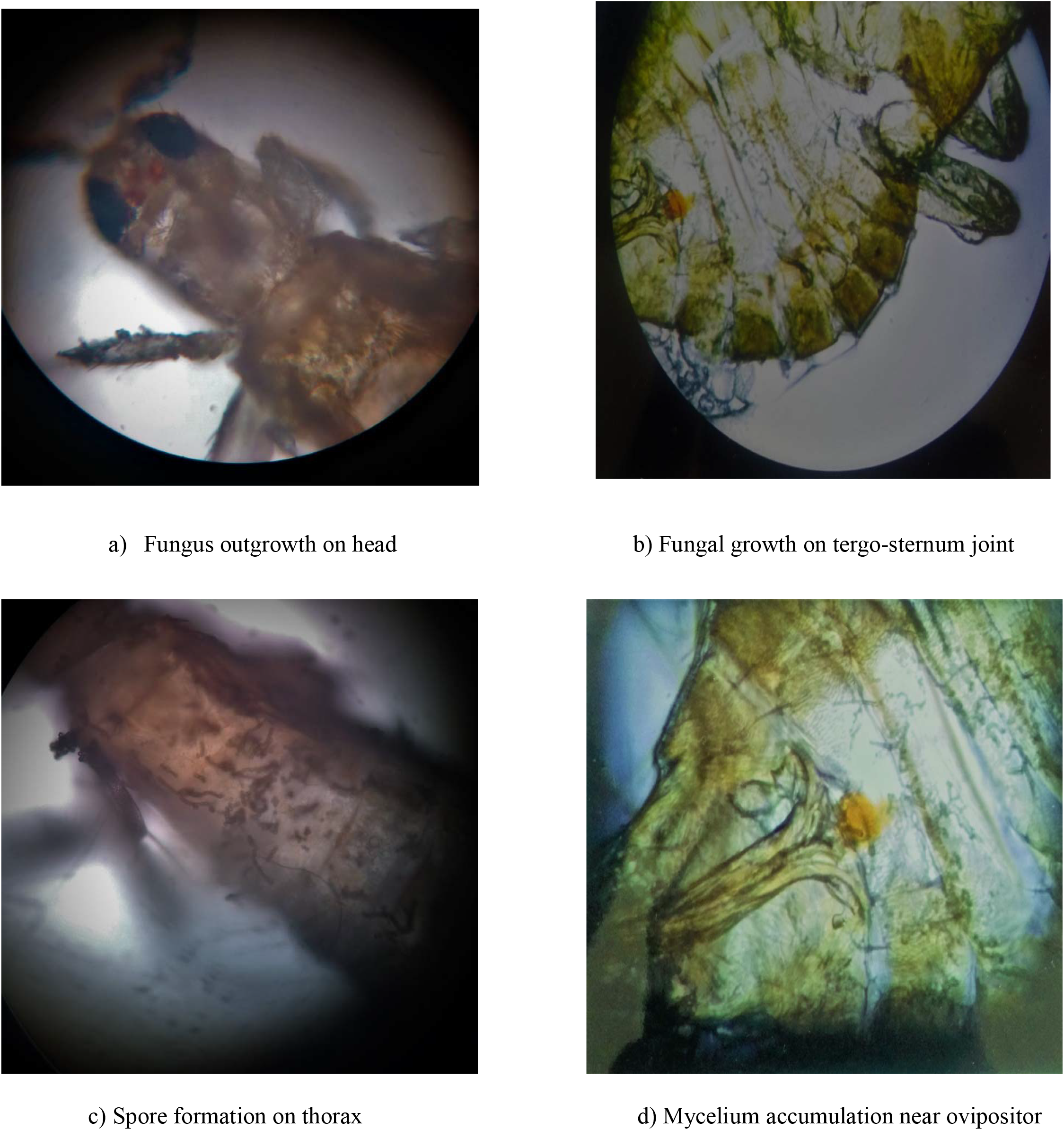
Mycosis of *Metarhizium anisopliae* on grape thrips.

**Plate 3:**
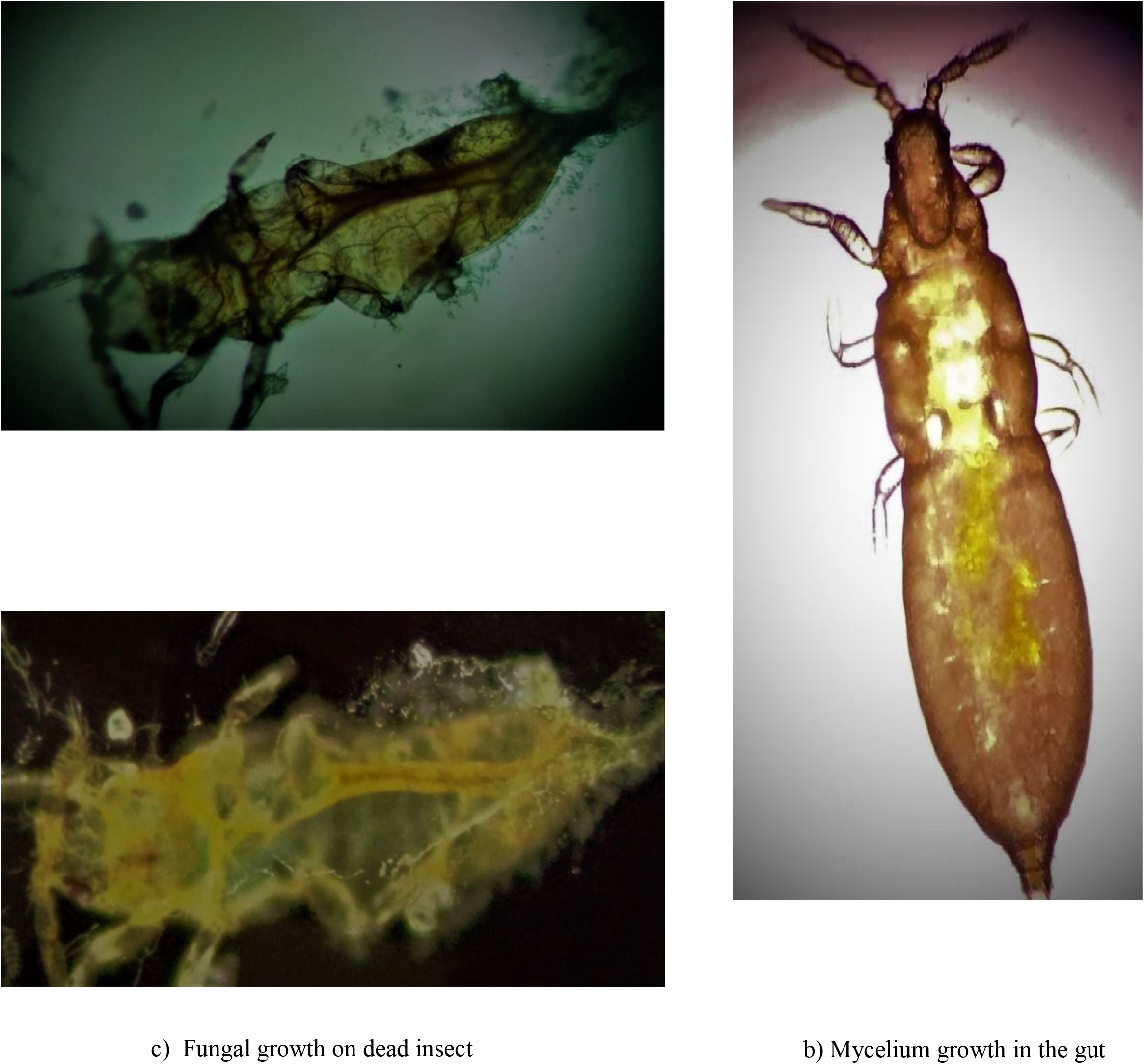
Mycosis of *Lecanicillium lecanii* on grape thrips.

## 4. Conclusion

Biorational insecticides neem oil 2% proved to be significantly superior over all other treatments, it was followed by entomopathogenic fungi like *Lecanicillium lecanii*, karanj oil 2%, *Metarhizium anisopliae* and *Beauveria bassiana* proved as best treatment in reduction of thrips population. Standard check emamectin benzoate 5% SG @ 11 g a.i.ha^−1^ proved to be significantly superior over all other treatments. The ICBR ratio in respect of different treatments ranged between 1.30 to 7.92. The highest ICBR ratio was observed among entomopathogenic fungi like *Lecanicillium lecanii* (1:6.34) and was followed by *Metarhizium anisopliae* (1:5.32), respectively. The cost benefit ratio of neem oil is (1:2.81). Mycosis by *Beauveria bassiana* was confirmed the pathogenicity of entomopathogenic fungi on grape thrips. Highly pronounced mycosis was observed by *Metarhizium anisopliae* on the dead bodies of thrips. Mycosis test of *Lecanicillium lecanii* was also proved on grape thrips.

## Acknowledgements

The authors gratefully acknowledge the technical support of Mahatma Phule Krishi Vidyapeeth, Rahuri, Dist. Ahmednagar

## References

Annual report. (2017-2018) National Research Centre for Grapes, Pune, Maharashtra, pp. 4–5.

Butani, D. K. (1979) Insects and Fruits, Periodical Export Book Agency, New Delhi, pp.190–194.

Dahiya, K. K., Lakra, R. K., and Ombir (1995) Studies on thrips infestation during reproductive stage of mango. Haryana Journal of Horticultural Science, 24:239–241.

Duraimurugan, P., and Jagadish, A. (2004) Control of *Scirtothrips dorsalis* Hood damaging rose flowers. Journal of Applied Zoological Researches, 15(2):149–152.

Flemming, R., and Retnakaran, A. (1985) Evaluation of single treatment data using Abbott’s formula with reference to insecticides. Journal of Economic Entomology 78(5):1179–1186

Kulakarni, N. S., Mani, M., and Banerjee, K. (2007) Management of thrips on grapes. National Research Centre for Grapes, Pune. Extension folder No. 13.

Kulakarni, N. S., Sawant, S. D., and Adsule P. G. (2008) Seasonal incidence of insect pests on grapevine its correlation with the weather parameters. Acta Horticulture, 785:313–320.

Latha, C. (2010) Evaluation of fungal pathogens against onion thrips, *Thrips tabaci* Lindeman (Thysanoptera:Thripidae). M Sc. (Agri.) Thesis, University of Agriculture Science, Bangalore.

Raikwar, M. K., Bhardvaj, V., Sawant, U., and Vora, V. (2011) Study of contamination level of pesticide residues in grapes (*Vitis vinifera*) in Maharashtra https://eco-web.com/edi/03336.html.

